# Automaticity in the reading circuitry

**DOI:** 10.1101/829937

**Authors:** Sung Jun Joo, Kambiz Tavabi, Sendy Caffarra, Jason D. Yeatman

**Author notes:** Correspondence should be addressed to Sung Jun Joo Department of Psychology, Pusan National University, 2, Busandaehak-ro 63beon-gil, Geumjeong-gu, Busan, South Korea.

## Abstract

Skilled reading requires years of practice associating visual symbols with speech sounds. Over the course of the learning process, this association becomes effortless and automatic. Here we test whether automatic activation of spoken-language circuits in response to visual words is a hallmark of skilled reading. Magnetoencephalography was used to measure word-selective responses under multiple cognitive tasks (N = 42, 7-12 years of age). Even when attention was drawn away from the words by performing an attention-demanding fixation task, strong word-selective responses were found in a language region (i.e., superior temporal gyrus) starting at ~300 ms after stimulus onset. Critically, this automatic word-selective response was indicative of reading skill: the magnitude of word-selective responses correlated with individual reading skill. Our results suggest that automatic recruitment of spoken-language circuits is a hallmark of skilled reading; with practice, reading becomes effortless as the brain learns to automatically translate letters into sounds and meaning.

## 1. Introduction

Mastering spoken language is natural, but learning a written language is not (Saffran et al., 2001; Wandell et al., 2012). Infants learn to understand spoken language through statistical regularities in natural speech starting from the earliest stages of development (Saffran et al., 1996). Indeed, encoding phonetic information during speech perception (Mesgarani et al., 2014; Yi et al., 2019) seems to be an automatic process in infants, and specialized circuits for processing spoken language located in the superior temporal gyrus (STG) are activated by speech sounds irrespective of attention, and even during sleep in infants as young as three months (Dehaene-lambertz et al., 2002). However, learning to read is an effortful process, which requires formal instruction on how to map arbitrary visual symbols (i.e., letters or graphemes) onto speech sounds (i.e., phonemes). Only after years of practice does this association become automatic and effortless, allowing for fluid and deep reading (Norton & Wolf, 2012; Wolf, 2018).

Behaviorally, the difference between a child who struggles to apply knowledge of grapheme-phoneme correspondence to decode a word and a child who fluidly reads a paragraph of text is striking. But neurally, what it means to automate the grapheme to phoneme conversion process is less clear. Cognitive models of reading have proposed that, for the literate brain, viewing printed words produces widespread and automatic activation of phonological and semantic representations (Harm & Seidenberg, 1999; Harm & Seidenberg, 2004; Seidenberg & McClelland, 1989; Van Orden & Goldinger, 1994). These models posit that literacy involves automatizing the connections between orthographic (visual), phonological, and semantic codes in the brain. Consistent with the prediction of these models, skilled adult readers show activation in canonical language processing areas such as the left inferior frontal gyrus (i.e., Broca’s area, IFG) and superior temporal gyrus (i.e., Wernicke’s area, STG) in response to visually-presented words regardless of whether or not the task requires them to actively read the words (Klein et al., 2015; Pattamadilok et al., 2017; Paulesu, 2001; Price, 2012; Turkeltaub et al., 2003; Wilson et al., 2004). Furthermore, there is ample behavioral evidence suggesting automatic involvement of phonological processing in response to printed words (Perfetti et al., 1988; Perfetti & Bell, 1991; Stroop, 1935). Indeed, in a series of studies examining the construction of “audiovisual objects” from text, Blomert and colleagues have suggested that automatization of letter-sound knowledge is a hallmark of skilled reading and the lack of automatization is a critical component of the struggles observed in children with developmental dyslexia (Blau et al., 2009; Blomert, 2011; van Atteveldt et al., 2004).

An intriguing conjecture is that neurons throughout the reading circuitry become automatically responsive to text, regardless of whether a subject intends to read the text, as a result of long-term simultaneous neural activity occurring in visual and language regions – akin to Hebbian learning – over the course of schooling (Hebb, 1949). Thus, becoming a skilled reader might involve automatizing the information transfer between visual and language circuits such that canonical speech processing regions in the STG start responding to written language even in the absence of attention.

In the present study, we defined automaticity as the evoked responses to visual stimuli in the absence of attention. To examine automaticity in the visual word recognition circuitry, we compare the response evoked by words to the response evoked by visually matched stimuli (scramble) under two different task conditions where attention is either focused on the stimuli (lexical decision task) or diverted away from the stimuli (color judgement on a fixation dot). Other studies have measured task effects within the reading circuitry (Chen et al., 2013; Chen et al., 2015; Mano et al., 2013). For example, Chen and colleagues (Chen et al., 2013) had subjects view words while performing either a (a) semantic judgement task, (b) lexical decision task, or (c) silent reading. They found that the cognitive task affected the evoked response to words as early as 150ms after stimulus onset indicating flexibility in the reading circuitry. In later work, they argued that the existence of task effects early in word processing is evidence against automaticity in word recognition (Chen et al., 2015). However, the question of automaticity need not be an either-or distinction: some computations in the reading circuitry might occur automatically while others might flexibly change based on the demands of the cognitive task (Kay & Yeatman, 2017). For example, a large body of studies have examined automatic audio-visual integration of visual symbols and speech sounds during letter processing (Brem et al., 2010; Raij et al., 2000; Taylor et al., 2019). Our concept of automaticity is distinct from these other studies; we examine the potential role of attention in gating information flow between visual and language cortex. We set out to ask whether neurons in the reading circuitry respond to visual word stimuli when visual attention is directed away from the stimuli. This is a classic manipulation used to dissociate bottom-up (task-independent) visual responses from top-down (task-dependent) responses in visual cortex (Fang et al., 2008; Kay & Yeatman, 2017).

Previous studies suggesting automaticity of word processing have not diverted attention from the stimuli. For example, although the Stroop task asks subjects to make an orthogonal judgment (color naming) rather than reading the word, attention is still directed toward the word stimuli (Strijkers et al., 2015; Stroop, 1935). The same is true for incidental reading tasks that direct attention to orthographic and shape features of the words (Klein et al., 2015; Pattamadilok et al., 2017; Paulesu, 2001; Price, 2012; Turkeltaub et al., 2003; Wilson et al., 2004).

To test our hypothesis, it is essential to disentangle bottom-up, visually-driven responses from top-down, task-related responses, and assess whether and how components of the reading circuitry are activated in an automatic manner by bottom-up signals from visual cortex. We used identical word stimuli in two tasks: one task is to read the word and decide whether the word is a made-up word (lexical decision task), and the other is to direct attention to the fixation mark and respond to rapid color changes (fixation task). By comparing responses to the identical stimuli in these two tasks, we could assess the extent to which word-selective responses require visual attention to words and whether the development of automaticity in the reading circuitry is related to children’s reading abilities.

We used magnetoencephalography (MEG) and source localization to define brain regions that were activated during a lexical decision task (active reading) and, within those regions, we characterized the time course of neural responses to text during a reading-irrelevant task in which words were placed outside the focus of attention. Using this paradigm, we first tested whether canonical speech processing regions show automatic responses to printed words. We then assessed whether the strength of automaticity in those regions depends on an individual’s reading skill.

## 2. Materials and methods

### 2.1. Participants

A total of 45 native English-speaking children ages 7-12 participated. We discarded data from 3 participants because their MEG signals were noisy and included data from the remaining 42 participants (age = 7.16 to 12.7 years, mean±sd = 9.6±1.5) for our analysis. Children without histories of neurological or sensory disorders were recruited from a database of volunteers in the Seattle area (University of Washington Reading & Dyslexia Research Database; http://ReadingAndDyslexia.com). Parents or legal guardians of all participants provided written informed consent under a protocol approved by the University of Washington Institutional Review Board. All participants reported normal or corrected-to-normal vision.

### 2.2. Reading ability assessment

Participants participated in a behavioral session in which they completed a series of behavioral tests. Reading scores were measured using the Test of Word Reading Efficiency (TOWRE-2), which measures the number of sight words (sight word efficiency, SWE) and pseudowords (phonemic decoding efficiency, PDE) read in 45 s. They also were assessed using subtests from the Woodcock-Johnson IV (WJ), which measures untimed sight word and pseudoword reading. Each test produces age-normed, standardized scores with a population mean of 100 and a standard deviation of 15. TOWRE and WJ measures of reading are highly correlated but also index slightly different aspects of skilled reading. The TOWRE measures the speed and automaticity of word recognition, while the WJ measures the ability to apply orthographic knowledge to decoding difficult words and pseudowords. Thus, for our study, we used TOWRE scores. We divided our participants into two groups: typical readers and struggling readers based on the TOWRE score of 80. This is a typical cut-point used to define children with dyslexia as it represents roughly the bottom 10% of the continuum. Table 1 shows the group comparison of age and reading scores between typical readers and struggling readers. Applying the same analysis using WJ scores did not change the pattern of the results. We also included subtests (verbal and matrix reasoning) from the Wechsler Abbreviated Scales of Intelligence as a general cognitive assessment.

**Table 1.** Behavioral test results between typical and struggling readers. Only reading related tests (TOWRE, IQ Vocabulary, Lexical task) showed group differences.

|  | Typical readers (n = 17) | Struggling readers (n = 25) | p-value (t-test) |
| --- | --- | --- | --- |
| Age | 9.25 (1.48) | 9.96 (1.49) | 0.15 |
| Reading skill (TOWRE) | 97.41 (13.58) | 67.24 (8.22) | 8x10<sup>-11</sup> |
| IQ Vocabulary | 57.65 (10.75) | 51.04 (7.59) | 0.03 |
| IQ Matrix Reasoning | 51.88 (8.75) | 48.96 (7.39) | 0.27 |
| Lexical task: d-primes | 1.74 (0.89) | 1.08 (0.88) | 0.03 |
| Fixation task: Reaction times (ms) | 520 (67) | 592 (152) | 0.10 |

### 2.3. Stimuli and experimental procedure

All the procedures were controlled by in-house python software (expyfun: https://github.com/LABSN/expyfun). Figure 1 shows the procedure of the experiment. The stimuli (real and pseudowords) were generated by manipulating the phase coherence of black word stimuli with a contrast of 25.4% on a gray background. Specifically, a Fourier transform of the image was computed, the phase component was shuffled, and a new image was generated by mixing a percentage of the scrambled image with the original image. The amount of phase scramble was 20% (clearly visible) and 80% (unreadable), corresponding to the word and scramble noise condition, respectively. We used phase scrambling of word stimuli to maintain low-level visual properties equivalent across the word (20% scramble) and scramble (80% scramble) conditions, so different responses to these conditions would be mediated by the process of reading but not by low-level visual processing. The stimuli were displayed on a gray background (50 cd/m2) of a back-projected screen using a PT-D7700U-K (Panasonic) projector. The stimuli subtended 2.7° at a viewing distance of 1.25 m.

**Figure 1.**
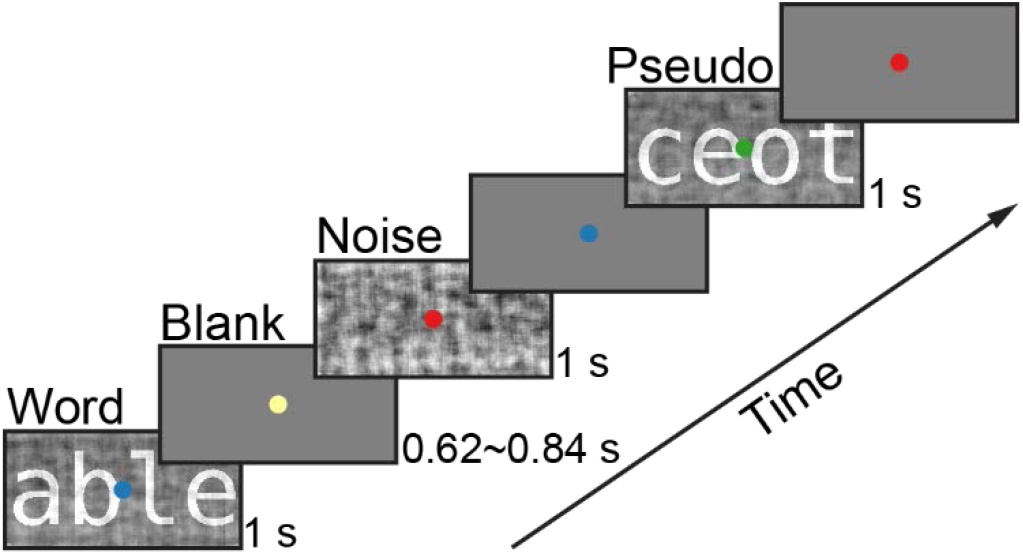
The experimental procedure. On a given trial, a word, a scramble noise, or a pseudoword was displayed for 1 s, followed by a gray blank screen. The next trial started after an inter-trial interval chosen randomly from a uniform distribution ranging from 0.62 to 0.84 seconds. Children conducted (1) a lexical decision task and (2) an attention-demanding fixation task in separate runs. The stimuli remained identical across both runs. During the lexical decision task, children were instructed to press a button when a pseudoword was presented. During the fixation task, they were instructed to press a button when the color of the fixation dot changed to red. Our analysis focused on a comparison of the response to words versus scramble since there were stimulus locked button-presses in response to the pseudowords that might interfere with MEG source localization.

We used MCWord (https://www.neuro.mcw.edu/mcword/) to select high-frequency four-letter words and generate orthographically plausible pseudowords matched in length (Supplementary Table 1). We created orthographically plausible pseudowords based on constrained trigram frequency.

On a given trial, the stimulus was displayed for 1 s followed by a blank screen with a random duration between 620 and 840 ms sampled from a uniform distribution. The fixation color changed every 500 ms during the stimulus presentation. The fixation color could be green, blue, yellow, cyan, or red. Participants conducted two tasks with this identical procedure (lexical decision task and fixation task) in separate runs (blocks). Thus, on each run, the visual stimuli were identical, but the task instructions changed. In the lexical decision task block, participants were asked to press a designated button when the word was a made-up word (pseudoword). In the fixation task block, they were instructed to press the button as quickly as possible when the fixation dot turned red (while ignoring the images). The lexical decision task allowed us to separately measure cortical responses that were automatically evoked by the stimuli (fixation task) as well as cortical responses that were associated with the cognitive task (reading). The experimental session had six experimental blocks (3 blocks for each task), and, in each block, there were 20 trials of the word condition, 20 trials of the scramble noise condition, and 7 trials of the pseudoword condition. The fixation task blocks (odd runs) and the lexical decision task blocks (even runs) were alternated in the session. Participants did one session, and there was a total of 60 trials for the word and the scramble condition and 28 trials for the pseudoword condition for each task. We did not counterbalance the order of tasks across participants because we aimed to study individual differences. Randomizing the order might cause some individual differences to be driven by experimental differences, which would be more problematic than the potential of incurring small biases in estimating the sample mean.

### 2.4. MEG and MRI data acquisition

MEG data were recorded inside a magnetically shielded room (IMEDCO) using a 306-channel dc-SQUID VectorView system (Elekta-Neuromag). Neuromagnetic data were sampled at 1kHz with a passband of 0.01 to 600 Hz. A 3D position monitoring system (Polhemus, Colhester, VT) was used to record the locations of head position indicator (HPI) coils, cardinal (nasion, left/right preauricular) anatomical landmarks. At least 100 digitized scalp points were used to coregister the MEG sensors with individual structural MRI. HPI coils were used to record the subject's head position continuously relative to the MEG sensors. Individual structural MRIs were obtained at The University of Washington Diagnostic Imaging Science Center (DISC) on a Philips Achieva 3T scanner for the boundary-element models that accurately characterize MEG forward field patterns. A whole-brain anatomical volume at 0.8×0.8 × 0.8mm resolution was acquired using a T1-weighted MPRAGE (magnetization prepared rapid gradient echo) sequence.

### 2.5. MEG data processing

The MEG data analysis including preprocessing and source localization were carried out using the MNE-Python and Freesurfer (http://surfer.nmr.mgh.harvard.edu/) dSPM pipeline according to community guidelines (Dale et al., 2000; Gramfort et al., 2013; Gross et al., 2013). Briefly, individual subject MEG data were first down-sampled (300 Hz), and denoised using signal space separation (Taulu et al., 2005) to remove environmental artifacts. Using the head position data from continuous HPI data from acquisition, the recorded MEG signal was compensated for head movements using the subjects initial head position as the target recording frame. Next the neuromagnetic data were low-pass-filtered (40 Hz cutoff), and signal space projection was used to suppress cardiac and ocular muscle artifacts identified using peripheral physiologic (ExG) sensor data. The resulting MEG signal data were windowed into 1000ms epochs of evoked neuromagnetic activity (including 100ms pre-stimulus baseline). Noisy trials were rejected based on peak-to-peak amplitude criteria for MEG magnetometer (30 pT) and gradiometer (4 pT/cm) channels. Evoked trial data were DC drift corrected using a mean baseline correction approach. Based on the number of trials in each condition after artifact rejection, we discarded data from 3 participants because there were not enough trials (< 40 trials in any of the conditions) for estimating average responses. Event related fields (ERFs) were obtained by averaging the remaining artifact-free trials of each participant and condition.

For individual subject MEG source reconstruction, an anatomically constrained three-compartment boundary element model (BEM) of the tesselated cortical surface was used to represent the cortical surface. As such, the pial surface was defined using a 3 mm grid consisting of 10242 dipoles per hemisphere. Dipole sources were constrained to be normal to the tangent at each location along the surface representing the pial boundary segmented from the structural MRI using Freesurfer watershed algorithm (http://surfer.nmr.mgh.harvard.edu/). The resulting individualized BEM was used to provide an accurate forward solution mapping dipole currents in the source space to the recorded ERFs. Using this colocation information and an iterative L2 minimum-norm linear estimator with shrunk noise covariance from the baseline sensor covariance (Engemann & Gramfort, 2015) we computed dynamic statistical parametric maps (dSPM) of conditional ERFs. For group level analysis individual dSPMs were mapped to an average brain (freesurfer averaged brain) using a non-linear spherical morphing procedure (20 smoothing steps) that optimally aligns individual sulcal–gyral patterns (Fischl et al., 1999).

To identify MEG sensors and vertices showing a word-selective response (words > scramble) during the lexical decision task (active reading), we ran a spatiotemporal cluster permutation t-test with no spatiotemporal constraints (Maris & Oostenveld, 2007). Note that we used this analysis only to define clusters of sensors and vertices and restrict our further analysis on them to assess word-selective responses during the fixation task.

For the sensor level analysis, we calculated pairwise t-statistics corrected for multiple comparisons using 2000 permutations and cluster level correction. We defined significant clusters (p < 0.001), and we averaged the sensors in the cluster to estimate the timecourse of MEG responses for each condition. The same sensors were used to estimate the timecourse of MEG responses in the fixation task. For the source level analysis, we calculated pairwise t-statistics corrected for multiple comparisons using 1024 permutations and cluster level correction. The resulting significant (p < 0.05) clusters in space and time were then visualized on the cortical surface. We further restricted vertices by selecting vertices that had a significant duration greater than 100 ms. We then used these regions that were localized during the lexical decision task runs to examine source activity during the fixation task runs (which were independently run in separate blocks). This region of interest (ROI) approach increases statistical power and limits the number of statistical comparisons used to test our main hypothesis while avoiding a possible circular analysis (Kriegeskorte et al., 2009).

## 3. Results

### 3.1. Behavioral results

Table A summarizes participants’ age, behavioral data, and IQ scores in typical and struggling readers. D’ for the lexical decision task suggest that all our participants performed the task as instructed (typical readers: 1.74±0.23; struggling readers: 1.08±0.17, mean±sem; p = 0.03, independent t-test; Bayes Factor B_10_ = 3.05). All word stimuli were four letter words with high lexical frequency to encourage our young participants, including struggling readers, to do the task. Despite the low performance compared to typical readers, the d-prime of struggling readers is above 1 suggesting that they also performed the task well. For the fixation task, typical readers and struggling readers performed the task equally well. The reaction times were 520±17 ms (typical readers) and 592±29 ms (struggling readers) (p = 0.15, independent t-test; Bayes Factor B_10_ = 0.76). The hit rates were 84±4 % (typical readers) and 73±4 % (struggling readers) (p = 0.10, independent t-test; Bayes Factor B_10_ = 1.02). Thus, differences in MEG responses during the fixation task could not be attributed to differences in performance between the two groups. Furthermore, each group’s non-verbal IQ scores were not different suggesting that any effects in the behavior and MEG responses were not due to differences in IQ.

### 3.2. Sensor-level analysis

We first ran an assumption-free spatiotemporal cluster analysis on the sensor data from the lexical task to find sensor clusters showing word-selective responses (words > scramble) during the epoch. We found a left-lateralized cluster, including temporal sensors (Figure 2(a)). Figure 2(b) shows the averaged time course on these sensors during the lexical task. The difference in response between words and scramble began at 243 ms after stimulus onset and lasted until 558 ms. This word-selective response was evident in typical readers (Figure 2(b), middle panel). The response to words was greater at [226, 396] ms compared to scramble. On the contrary, in struggling readers, the same sensors showed less clear word-selective responses (Figure 2(b), bottom panel; [310, 376] ms).

**Figure 2.**
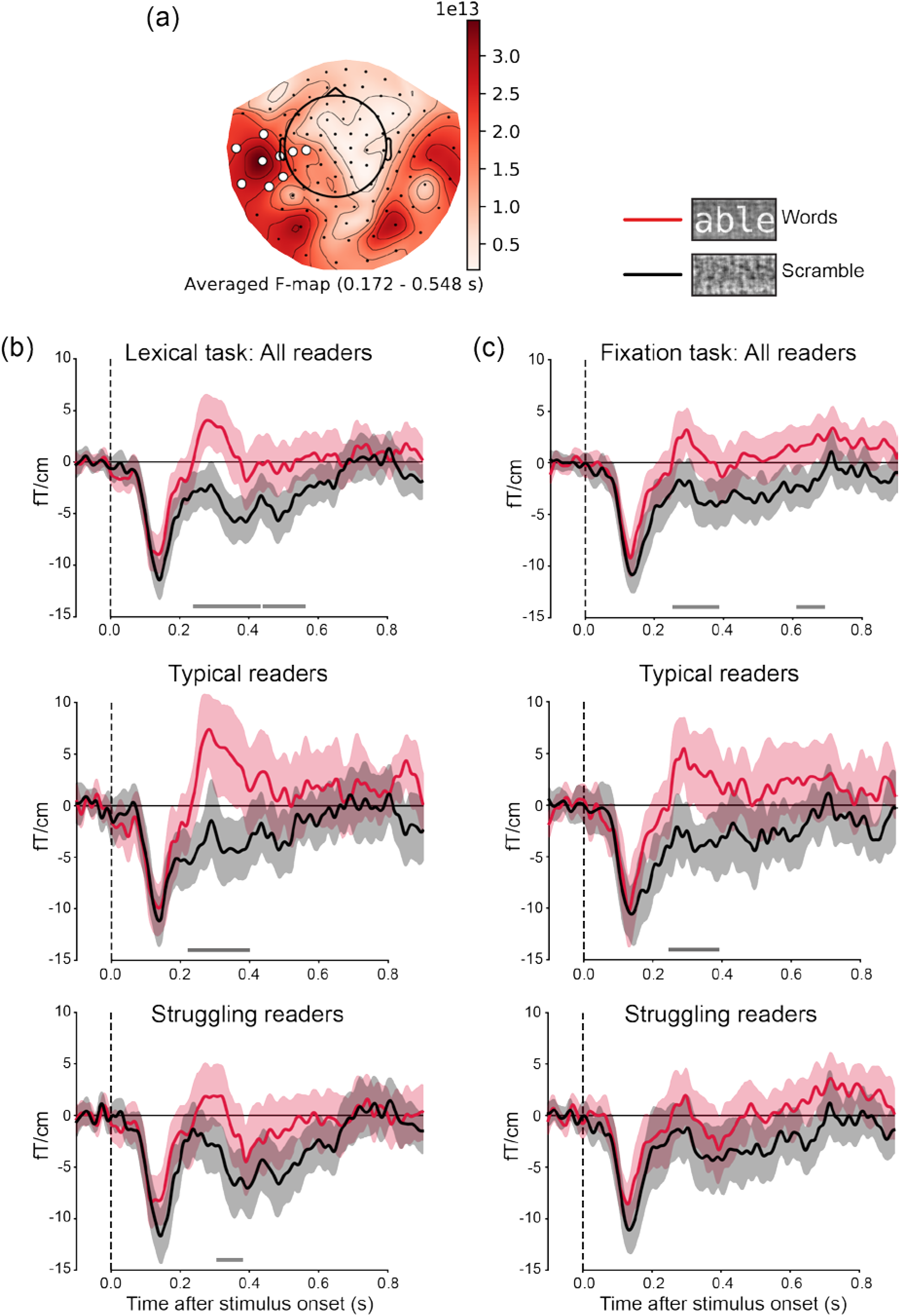
Sensor level word-selective responses. (a) Spatiotemporal cluster-based permutations resulted in significant clusters showing greater responses to words compared to scramble in the lexical decision task. White dots represent sensors in the clusters and are averaged to produce MEG time courses. The color bar indicates F-values. (b) The MEG time course in the lexical decision task for all participants (top), typical (middle), and struggling readers (bottom). (c) The MEG time course in the fixation task in the same clusters defined using lexical decision task data for all participants (top), typical (middle), and struggling readers (bottom). The red and black lines represent the response to words and to scramble, respectively. The shaded areas are 68% confidence intervals equivalent to ±1 sem. The gray bars at the bottom of each graph indicate significant time points from the permutation t-test using the sensors in the cluster (p < 0.001).

To test whether the same sensors showing word-selective responses during the lexical decision task also show word-selective responses during the fixation task, we used the same sensors to characterize the time course in the fixation task. We found word-selective responses at [258, 383] ms in those sensor clusters even though participants’ attention was directed away from the visual word stimuli (Figure 2(c), top panel). Critically, this word-selective response was only present in typical readers (Figure 2(c), middle panel; [250, 388] ms), but not in struggling readers (Figure 2(c), bottom panel).

To assess the relationship between reading skill and word-selective responses in this cluster, we averaged the time course within a time window of [283, 383] ms in both the lexical decision and the fixation task (note the cluster was defined orthogonally to this analysis). We found that word-selective responses in the lexical decision task were correlated with reading skill (r = 0.39, p = 0.009; Figure 3(a)). We further confirmed this correlation by calculating the skipped correlation (r = 0.33, CI_95%_ = [0.01, 0.57]), which estimates more robust correlation between variables by detecting and removing outliers (Pernet et al., 2013). In the fixation task, the correlation between word-selective responses and reading skill was similar to the one in the lexical decision task although the effect size is smaller (r = 0.31, p = 0.048; skipped correlation r = 0.31, CI_95%_ = [−0.003, 0.559]; Figure 3(b)).

**Figure 3.**
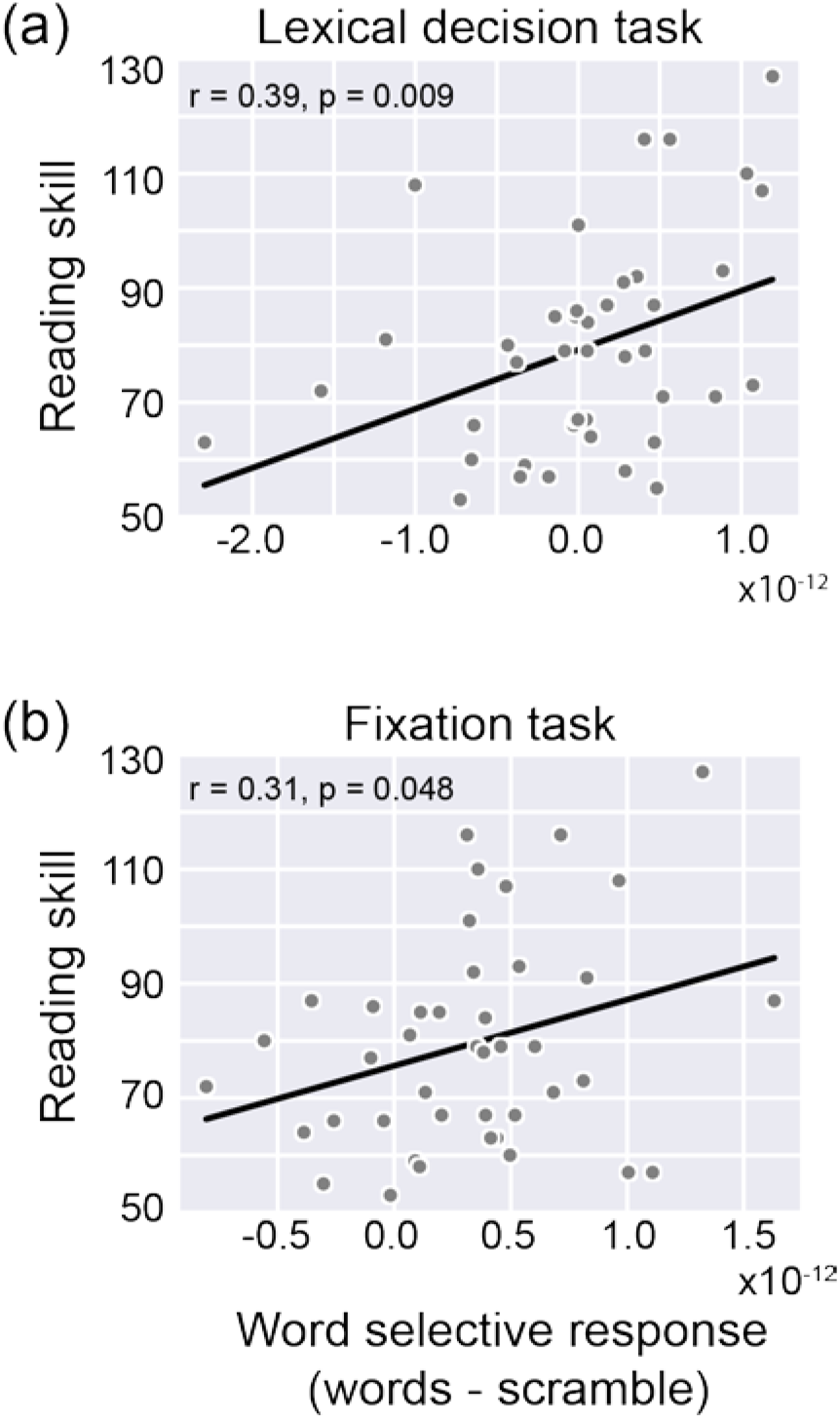
The correlation between the word-selective response and reading skill at the sensor level. (a) The word-selective response and reading skill is correlated in the lexical decision task. (b) In the fixation task, the correlation is similar to the correlation in the lexical decision task. The y-axis is the standardized reading score (mean = 100, sd = 15) and the x-axis is the word-selective MEG response (word-scramble).

These results suggest that there may be automatic word-selective responses during the fixation task in typical readers even when they are not paying attention to the words. To understand the neural sources that contribute to these effects, we performed a source localization analysis.

### 3.3. Lexical decision making activates language processing network

To assess automaticity in the reading circuitry, we localized cortical regions of interest (ROIs) that were engaged during the lexical decision task, and then assessed the timing and magnitude of visually evoked responses to text during the fixation task. This allowed us to dissociate visual, bottom-up responses to printed words from active reading related responses. To correct for multiple comparisons in both space (20,484 vertices) and time (301 time points), we employed the conservative spatiotemporal clustering algorithm (Maris & Oostenveld, 2007): ROIs were defined as significant vertices resulting from a permutation t-test (2-tailed) between the word and scramble noise conditions in the lexical decision task (word > scramble, Figure 4).

**Figure 4.**
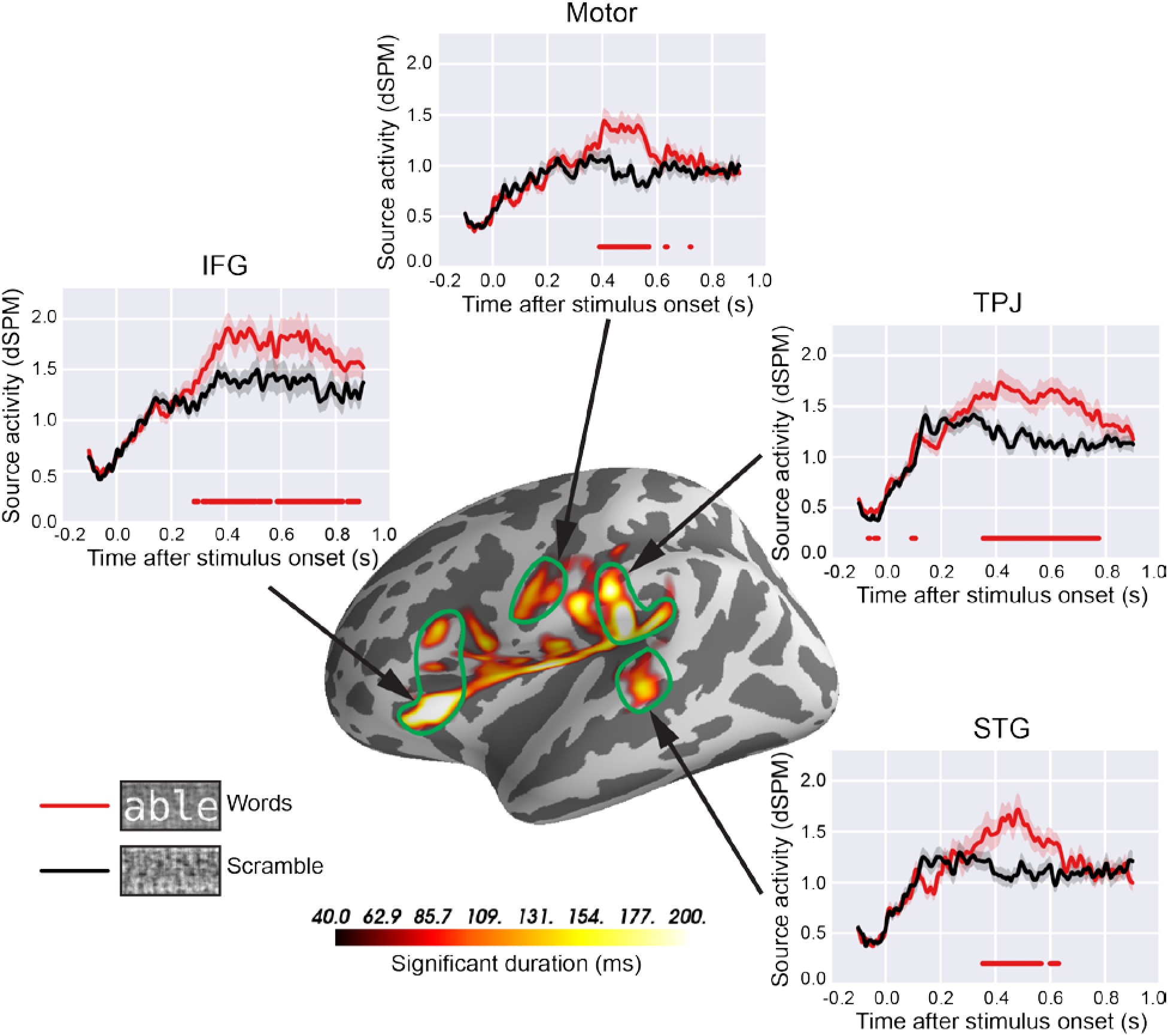
Canonical language regions respond to printed words during the lexical decision task. Cortical activation map in the left hemisphere during the lexical decision task projected into the freesurfer averaged cortical surface template (fsaverage), and thresholded based on spatial temporal clustering to correct for multiple comparisons. The color bar represents the duration of time that each vertex shows a significantly greater response to words compared to scramble after correcting for multiple comparisons. The red and black lines show the source activity for words and scramble, respectively. The red dots at the bottom of the plot indicate timepoints where the response is significantly different between the two conditions (determined by bootstrapping analysis p < 0.05).

Figure 4 shows the spatiotemporal extent of significant neural activity to words compared to scramble in the left hemisphere during the lexical decision task. Each inset shows the time course of individual ROIs computed using dynamic statistical parametric mapping (dSPM) (Dale et al., 2000), averaged across all subjects for word (red) and scramble (black) conditions. Consistent with previous literature, we found that canonical language processing regions are engaged while subjects performed a lexical decision task on visually presented words (Helenius et al., 1998). We defined four main ROIs based on the significant vertices in this cortical activation map (word > scramble): left inferior frontal gyrus (IFG), left temporoparietal junction (TPJ), and left superior temporal gyrus (STG) and left sensorimotor cortex. The posterior regions (STG and TPJ) are the conventional loci associated with phonological processing and are activated across auditory speech perception and reading tasks (Pugh et al., 2001). The IFG (Broca’s area) is associated with different components of linguistic functions such as semantic, lexical, and phonological processing in both spoken and written languages (Sahin et al., 2009).

The difference in responses to words (red) and scramble (black) began to diverge around 350 ms after stimulus onset in posterior ROIs (Figure 4 insets; STG: 353 ms, TPJ: 356 ms, motor: 390 ms, bootstrapping analysis p < 0.05). In contrast, there was earlier neural activity in IFG starting at ~280 ms, consistent with previous evidence for early MEG source synchronization and electrocorticography recordings during lexical processing (Cornelissen et al., 2009; Klein et al., 2015; Sahin et al., 2009; Wheat et al., 2010).

Figure 5 shows the time course of MEG source estimates in typical and struggling readers. To characterize neural activity associated with the lexical decision task, we calculated the average differences in MEG source estimates between the word and scramble condition during the 100 ms temporal window of [350, 550] ms showing highest word-selective responses (word-scramble) across ROIs (Supplementary Figure 1).

**Figure 5.**
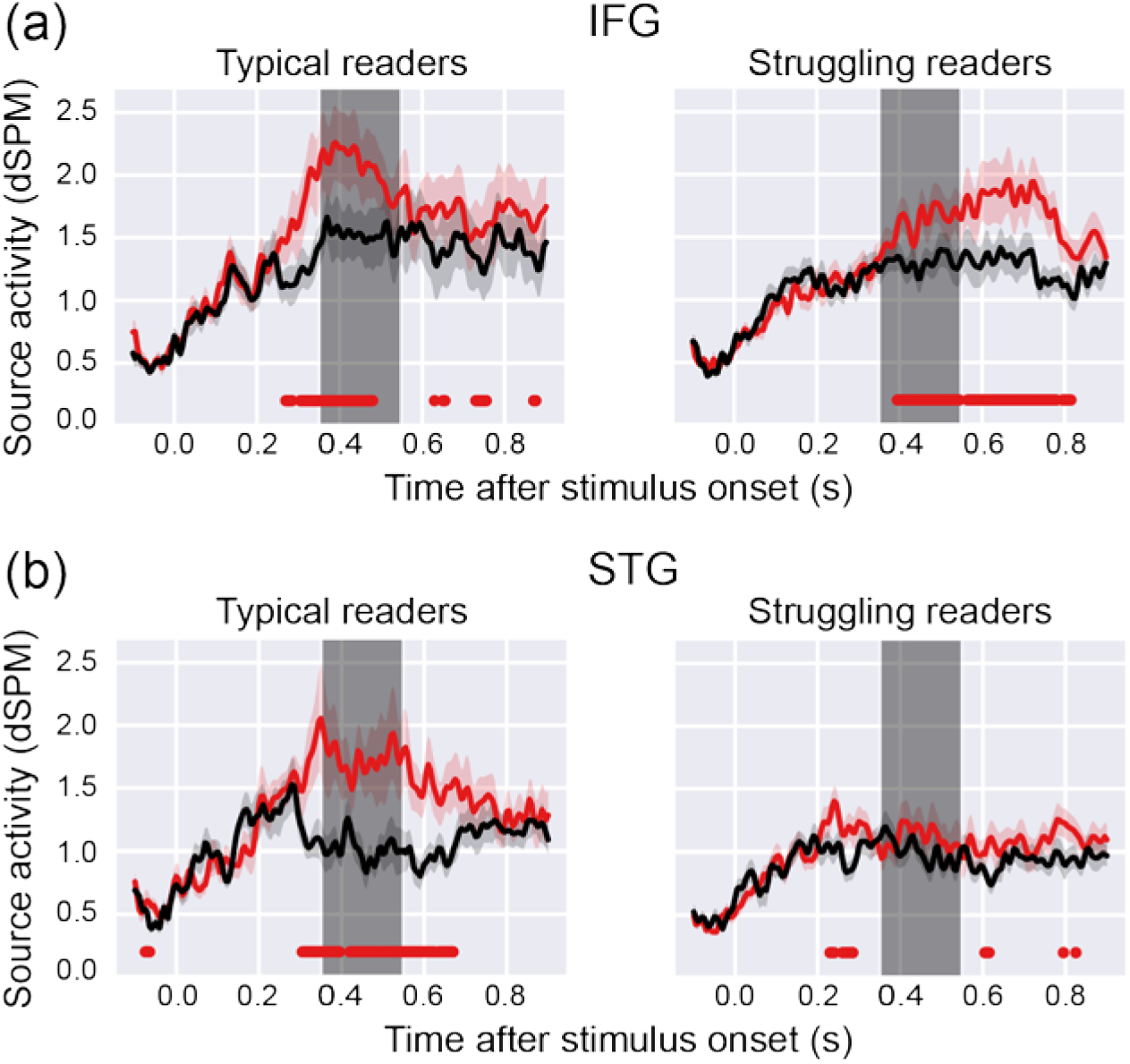
Cortical activity during the lexical decision task correlates with reading skill. Responses within each functionally-defined region of interest are shown separately for typical versus struggling readers in the (a) Inferior Frontal Gyrus (IFG) and (b) Superior Temporal Gyrus (STG). In the IFG, there is larger response to words compared to scramble for both typical and struggling readers. In the STG there is a robust difference in the response to words versus scramble only for typical readers. In struggling readers, STG activity does not differ for the two stimulus types. The red and black lines show the source activity for words and scramble, respectively. The shaded areas represent ±1 s.e.m. across participants. Red dots represent time points showing significantly larger responses to words compared to scramble (determined by bootstrapping analysis p < 0.05). The gray shaded rectangles are the time windows used to calculate the averaged response. (c) Only the STG response correlates with reading skill. The y-axis is the standardized reading score (mean = 100, sd = 15) and the x-axis is the word-selective source activity (word-scramble).

In the IFG, struggling readers showed a slower response to words that diverged from the response to scramble at 400 ms and was sustained until 800 ms post-stimulus onset (Figure 5(a)). In comparison, typical readers showed a markedly faster and shorter enhanced response to words beginning at 277 ms and lasting up to 500 ms after stimulus onset. The word-selective source activity in the IFG (word-scramble) in the time window of [350, 550] ms post-stimulus interval of interest did not correlate with reading skill (r = 0.25, p = 0.11) suggesting that task-related activity in the IFG does not differ substantially between good versus poor readers.

In the STG (Figure 5(b)) and TPJ, there were robust and highly significant responses to words compared to scramble in typical readers, while struggling readers showed much weaker responses. Within these regions, the word-selective source activity was correlated with reading skill (STG: r = 0.50, p = 0.0006, Figure 5(c); TPJ: r = 0.46, p = 0.002, data not shown). We further confirmed this correlation by skipped correlation (STG: r = 0.35, CI95% = [0.03, 0.63]). These results suggest different roles for each of these regions in the lexical decision task: the IFG was engaged in the lexical decision task early and irrespective of reading skill, while word-selective neural activity in the STG and TPJ depended on reading skill. Next, we capitalize on the independent data set obtained during the fixation task to (a) examine which of these responses to text are task-dependent and (b) whether the correlation with reading skill depends on the task performed by the subject or reflects the strength of the bottom-up response to text.

### 3.3. Automatic responses to text in speech processing regions

Next, we tested the hypothesis that, for skilled readers, there is an automatic, bottom-up response to visually presented words in language regions, even in the absence of attention or conscious reading. We reasoned that if word-selective neural activity in the regions identified in the lexical decision task is equivalent during the fixation task, then it is evidence for an automatic, stimulus-driven, response. Based on results from the lexical decision task, we predicted that the IFG would show task-dependent responses to printed words whereas the STG and TPJ would show responses to printed words even in the absences of attention, likely indicative of the automatic association between graphemes and phonemes.

We found that the left STG showed highly significant responses to words compared to noise during the fixation task and the strength of the response depended on reading skill. Other areas showed no difference in responses to word and noise stimuli (Figure 6(a)). Only in typical readers, MEG responses to words (red) were greater compared to scramble (black), and the difference between the two diverged earlier (at ~300 ms) compared to the lexical decision task (Figure 6(b)). Importantly, as in the lexical decision task, the word-selective source activity in the [350, 550] ms post-stimulus interval (see Supplementary Figure 1) strongly correlated with reading skills (Figure 6(c); r = 0.48, p = 0.001, skipped correlation: r = 0.31, CI_95%_ = [0.02, 0.54]). In fact, individual word-selective source activity in the lexical decision and fixation tasks were highly correlated (Figure 6(d); r = 0.67, p = 1×10^−6^; skipped correlation: r = 0.33, CI_95%_ = [0.06, 0.57]). Furthermore, the word-selective responses in these two tasks were statistically equivalent across the entire epoch (Supplementary Figure 2). These results suggest that the left STG responded similarly to words over scramble regardless of whether subjects were actively reading them and performing a lexical decision task, or engaging in an attention-demanding task on the fixation dot (Figure 6(d)).

**Figure 6.**
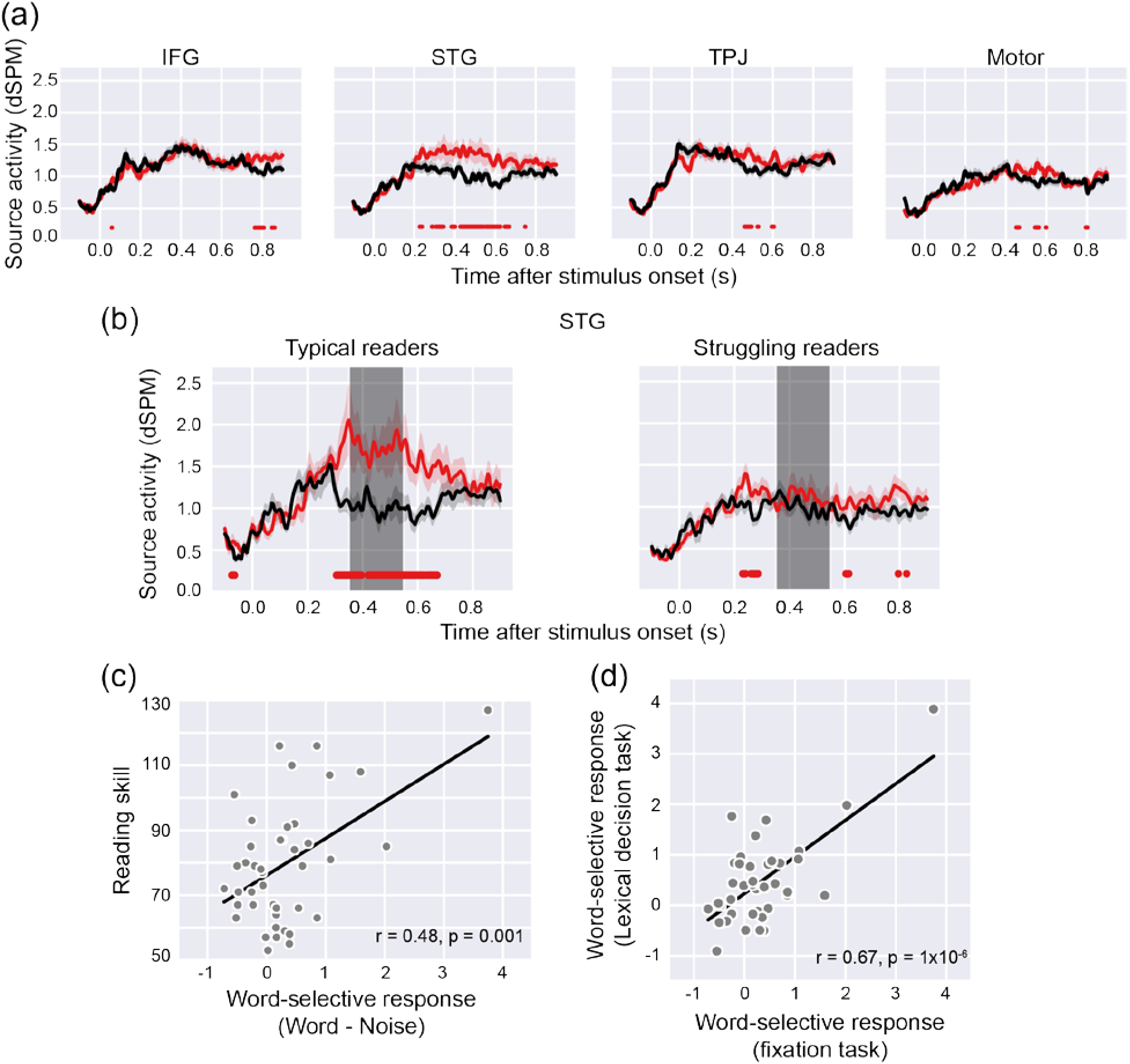
Automaticity in the superior temporal gyrus is related to reading skill. (a) The response to words (red) and scramble (black) for the full sample (N=42) during the fixation task is shown within the four regions that were localized independently with data from the lexical decision task (see Figure 4). At the level of the full sample, only the superior temporal gyrus (STG, second plot) shows a greater response to words versus scramble when attention is diverted from the stimuli. (b) This “automatic” response in the STG is driven by the children with relatively strong reading skills and is not present in the struggling readers. Gray shaded rectangles are the time window used to calculate the averaged source activity. (c) Word-selective source activity in the STG during the fixation task correlates with reading skill. The y-axis is the standardized reading score (mean = 100, sd = 15) and the x-axis is word-selective source activity (word-scramble). (d) Individual word-selective source activity in the STG during the fixation task (x-axis) are highly correlated with word-selective source activity during the lexical decision task (y-axis).

Thus, even though the STG response might be modulated by task demands in some cases, a subject showing a weak word-selective response on the fixation task still shows a weak word-selective response when prompted to actively attend to and analyze the text. This finding underscores the importance of automaticity for skilled reading; akin to the visual system where attention and task demands modulate the automatic bottom-up response of neurons that code specific features of the visual stimulus, task demands might serve to modulate the automatic bottom-up response in the STG but only to the extent that the STG is responsive to text.

Although the left TPJ showed greater responses to words than scramble during the lexical decision task, we did not find a similar pattern during the fixation task. Typical readers showed some separation in responses to words compared to scramble, but the difference was much smaller than in the left STG. There was a trend, but no significant correlation between word-selective source activity and reading skill (r = 0.29, p = 0.07). Furthermore, in the left IFG, there was an equivalent response to words and scramble stimuli during the fixation task, and this response did not differ by group or relate to reading skill. To ensure that the absence of a word-selective response in the IFG during the fixation task did not reflect the specific ROI, we confirmed this finding based on spatiotemporal clustering of the fixation task data: no frontal lobe clusters were revealed in the word > scramble contrast. Together, these results suggest that the left STG is automatically engaged in response to visually presented words, and that the strength of neural activity is associated with reading skills in children. The left IFG, on the other hand, is important for carrying out the lexical decision task, but is not critically involved in automatic grapheme-phoneme conversion processing.

### 3.3. Reading disability versus reading level

So far we have shown that, in strong readers, there are robust and automatic responses to text in canonical speech processing regions (i.e., left STG). Struggling readers show little or no activation in the left STG on the fixation or lexical decision tasks. Since our sample included children of different ages (between first and fourth grade) and, across those ages, we include both typical and struggling readers, we next seek to determine the relative contributions of age versus reading ability to the STG response. In other words, is the lack of left STG response indicative of a deficit in the circuit in children with reading difficulty? Or, does automaticity steadily increase over each year as children's reading skills improve? To answer these questions, we first examined the correlation between age and the left STG response. We found that the left STG response to words did not increase with age (fixation task: r = 0.06, p = 69; lexical task: r = 0.19, p = 0.24). Next, we used a multivariate regression model to test the additive contributions of age and reading ability (indexed by age-normed scores on the TOWRE) to the left STG response. We found a highly significant relationship between reading ability and left STG response, irrespective of age and, once again, found no contribution of age in the model (Table 2). Thus, automaticity in the left STG response to text is likely to develop early in literacy learning for typically reading children, establishing a foundation for children to hone their reading accuracy, rate and fluency. For struggling readers, the lack of response in the left STG is likely to represent a barrier that continues to affect their ability to learn reading skills. In line with this perspective, every measure of reading skills including real and pseudoword reading accuracy and speed and fluency all correlated with the left STG after controlling for the age of the subjects (Table 2).

**Table 2.** Superior temporal gyrus response is related to reading skills but not age. Coefficients from regression models examining the relationship between scores on the Test of Word Reading Efficiency (TOWRE, Model 1), Woodcock Johnson Basic Reading Skills Composite (BRS, Model 2), Woodcock Johnson Reading Fluency (RF, Model 3) and the word-selective STG response (words - scramble), controlling for age.

|  | Coefficient | Std. error | p-value |
| --- | --- | --- | --- |
| **Model 1** |  |  |  |
| Age | 0.077 | 0.07 | 0.293 |
| TOWRE | 0.020 | 0.01 | 0.002 |
| **Model 2** |  |  |  |
| Age | 0.076 | 0.07 | 0.311 |
| BRS | 0.019 | 0.01 | 0.006 |
| **Model 3** |  |  |  |
| Age | 0.062 | 0.08 | 0.412 |
| RF | 0.014 | 0.01 | 0.015 |

## 4. Discussion

We have demonstrated that a part of canonical circuitry for processing spoken language, the left superior temporal gyrus (STG), is automatically engaged when skilled readers view text. It has long been hypothesized that automating the association between printed symbols and spoken language is at the foundation of skilled reading (Blau et al., 2009, 2010; Blomert, 2011; Harm & Seidenberg, 1999; Seidenberg & McClelland, 1989; van Atteveldt et al., 2004) and our results formalize this concept of automaticity and its relationship to reading skill: In skilled readers, text evokes a word-selective response in the STG even when subjects are performing a distracting task with attention directed away from the word stimuli. Furthermore, the magnitude of the STG response is strongly correlated with reading skill and largely consistent across both the fixation and the lexical decision task. In contrast, neural activity in the IFG depends on the task. The IFG shows word-selective responses during the lexical decision task but not during the fixation task. While there is always the possibility that this task effect in the IFG reflects either (a) task order or (b) the specific ROI definition, there is a striking difference between the response profiles in the IFG versus the STG. One interpretation of this dissociation is that the STG response on the fixation task is a bottom-up, automatic, stimulus-driven response while the IFG response is associated with the specific task the subject performs while viewing a word.

Our results provide evidence of automatic responses to printed words in language processing regions by dissociating bottom-up responses from task-related responses. This inference is not at odds with data demonstrating that task demands modulate responses throughout the reading circuitry (Chen et al., 2013; Chen et al., 2015; Kay & Yeatman, 2017); future work should systematically catalogue the effects of task and stimulus manipulations. Our results are right in line with the prediction made by classic cognitive models of the reading architecture (Harm and Seidenberg, 2004; Pugh et al., 2001; Pugh et al., 2010; Seidenberg and McClelland, 1989). Furthermore, our findings are compatible with the implications of the word superiority effect (Reicher, 1969; Wheeler, 1970) in the sense that words have a privileged access to mental lexicon and this special representational status can reach high levels of automaticity. However, the neural underpinnings of the word superiority effect are still debated (Heilbron et al., 2020).

In previous experiments attempting to measure the automatic response to printed words, attention was still directed to visual stimuli while subjects were engaged in tasks other than reading (i.e., visual feature detection on word stimuli), making word stimuli task-irrelevant but still inside the focus of attention (Brunswick et al., 1999; Paulesu, 2001; Price et al., 1996; Turkeltaub et al., 2003). Thus, it is difficult to disambiguate the extent to which attention to the words provoked unwanted reading. In fact, in these previous experiments words activated brain regions (e.g., the IFG) that our data indicate are only active during reading tasks (e.g., lexical decision). By diverting attention from printed words, we show a dissociation between the IFG and STG response: while both areas are robustly activated during the lexical task, automatic responses to printed words (in the fixation task) are only found in the STG.

We used word-selective responses (words – scramble) as a proxy for reading-related activity in our study. The comparison between words and matched visual stimuli (phase scrambled noise patches) is widely used to define word-selective regions and reading skill dependent brain activity (Ben-Shachar et al., 2011; Caffarra et al., 2017; Kay & Yeatman, 2017; Lerma-Usabiaga et al., 2018; Rauschecker et al., 2012; Rauschecker et al., 2011; Tarkiainen et al., 1999). However, the contrast between word and scramble does not isolate specific aspects of word processing, and it is possible that the difference in responses to words and scramble might be caused by factors other than reading. For example, words are more familiar and meaningful stimuli than scrambled noise patches. There are several points to make regarding this issue. First, our results show that conventional language processing regions do respond more to words than matched visual stimuli. There is no literature showing attention gain to visual stimuli is restricted to brain regions with specific higher-level function. Rather, attentional effects can be found as early as primary visual cortex (Buracas & Boynton, 2007). Second, we observed word-selective responses in the fixation task when participants’ attentional state was equated between words and scramble. Lastly, only the left STG showed word-selective responses in the fixation task among many regions showing word-selective responses in the lexical decision task. Thus, our results cannot be attributed to simple visual familiarity effects.

Our findings stand in contrast to previous work in which words were rendered invisible due to rapid visual backward masking. In the case of masking, words do not elicit measurable functional magnetic resonance imaging (fMRI) responses in language regions despite clear behavioral priming effects (Dehaene et al., 2001). The discrepancy between our results and previously reported priming effects might be due to the difference between fMRI and MEG measurements. MEG might be more sensitive to brief neural activity compared to fMRI because of the sluggish nature of fMRI responses. Indeed, neural activity related to behavioral priming effects was found using EEG measurements (Luck et al., 1996), which have similar temporal resolution as MEG.

Another possibility is that the connectivity between visual and language circuits depends on the visibility of words. In the work by Dehaene and colleagues (Dehaene et al., 2001), a briefly presented word (30 ms) was rendered invisible due to visual backward masking. In contrast, words (83 ms) in the Luck et al.’s experiment were not invisible although the word detectability was reduced in the attentional blink paradigm (Luck et al., 1996). In our experiment, words were displayed for 1 s and had a high contrast, making them visible although attention was removed from them. It is an interesting future direction to study how the visibility of words affects automaticity in the language processing areas by manipulating noise levels (Ben-Shachar et al., 2011) and timing of the stimuli as well as task-demands (Kay & Yeatman, 2017).

In canonical perisylvian language processing regions, only the STG but not the TPJ (which is near proximity of supramarginal gyrus), showed automaticity. Many authors consider these two regions to have a similar function for reading and traditional “Wernicke’s area” is often presumed to include both regions. While both regions are associated with phonological processing during auditory word processing (Binder et al., 1994; DeWitt & Rauschecker, 2012), the supramarginal gyrus might be involved in further cognitive processing beyond phonological encoding. For example the supramarginal gyrus (and TPJ) might be involved in storing information for conducting tasks (Paulesu et al., 1993; Warrington et al., 1971) whereas neural activity in the STG is often found to occur independently of the cognitive task (Binder et al., 1994, 1997; Wise et al., 1991).

In the word superiority effect (Reicher, 1969; Wheeler, 1970), a letter is more detectable when the letter is embedded in a word than in a pseudoword. A recent study showed that neural representation of letters is enhanced when the letter is embedded in a real word compared to a pseudoword (Heilbron et al., 2020). Interestingly, they found functional coupling between the IFG and the posterior middle temporal gyrus, which is in close proximity to our STG ROI. In our study, word-selective responses during the lexical decision task begin early in the IFG compared to the STG (Figure 5). This is in line with previous studies showing early responses to words in the IFG (Cornelissen et al., 2009; Klein et al., 2015; Sahin et al., 2009; Wheat et al., 2010). It would be an interesting future direction to study whether and how connectivity between the IFG and the STG can predict reading skill dependent STG responses by combining diffusion MRI and MEG source localization methods (Bedo et al., 2014).

Based on our results, we can formulate a hypothesis that there might be similar neural activity in response to printed words as to auditory word stimuli in the STG. Future work should test whether automatic responses to printed words in the STG share similar neural codes to the responses to auditory words.

Overall, our study demonstrated automatic responses to printed words in the STG, a part of canonical language processing areas. Skilled reading seems to require coactivation in the reading network for spoken and written language (McCandliss et al., 2003; Price, 2012; Pugh et al., 2010). Interestingly, the level of coactivation in the left hemisphere reading network in early readers could be a predictor of reading outcomes after two years (Preston et al., 2016), suggesting that becoming a skilled reader relies on shared neural responses to both print and speech. Our results, showing the magnitude of automatic responses depends on reading skill, suggests that automaticity in the STG may be a hallmark of skilled reading; with practice, reading becomes effortless as the brain learns to automatically translate letters into sounds and meaning.

## Acknowledgements

We would like to thank Samu Taulu, Eric Larson, Patricia Kuhl and the rest of the I-LABS MEG team for helpful discussion and input throughout this project. This work was supported under the framework of international cooperation program managed by the National Research Foundation of Korea (NRF) (No. 2018K2A9A2A20088926 and 2019R1C1C1009383) to SJJ. This work was also funded by NSF BCS 1551330, NICHD R21HD092771 and R01HD09586101and Jacobs Foundation Research Fellowship to JDY. SC was funded by the European Commission (H2020-MSCA-IF-2018-837228-ENGRAVING).

---

**Supplementary Table 1.**
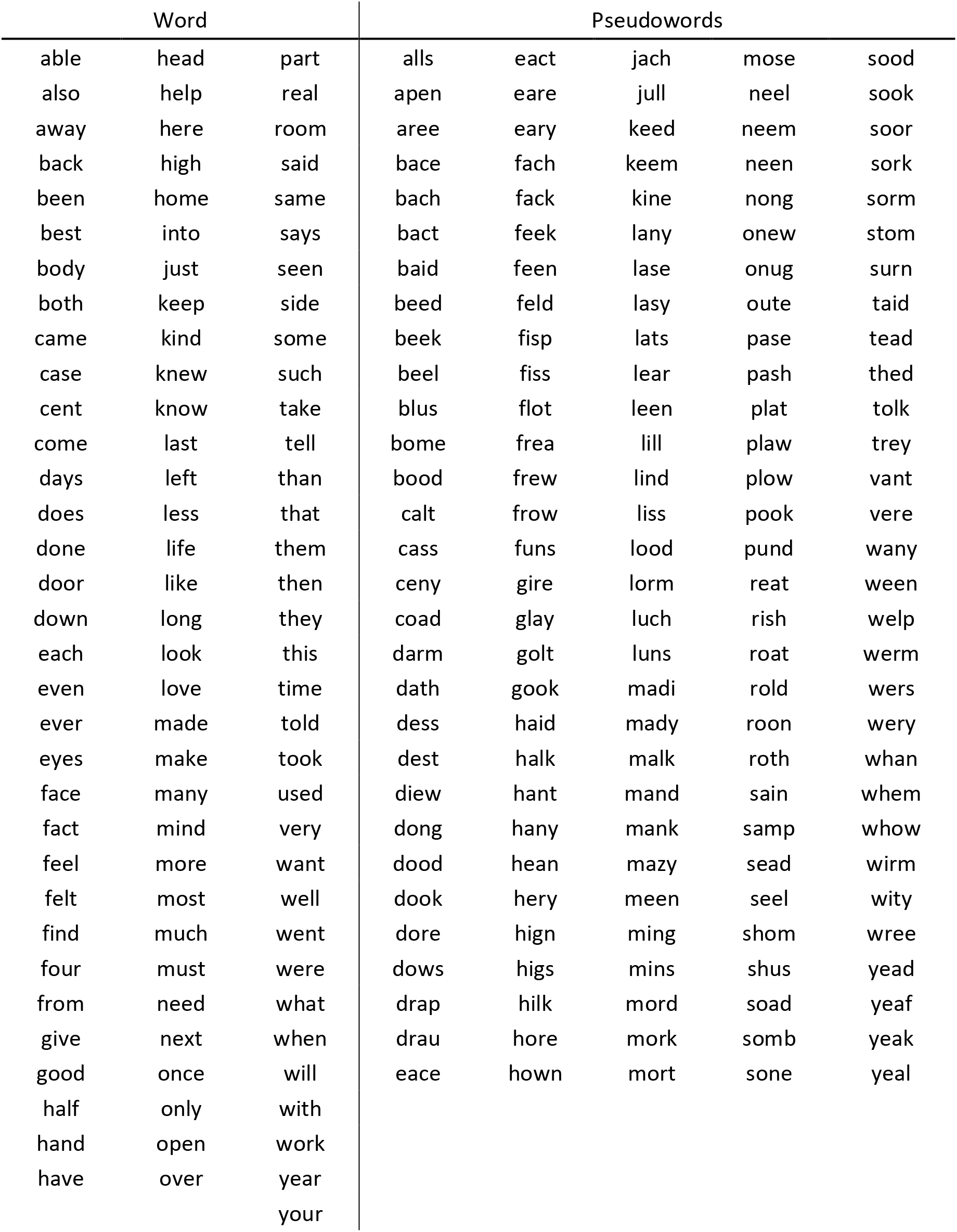
The word and pseudoword list used in the experiment. We used MCWord (https://www.neuro.mcw.edu/mcword/) to create both the word and the pseudoword list. We created 100 4-letter words with frequency of 300. We created orthographically plausible pseudowords based on constrained trigram frequency with the same length as the real words.

**Supplementary Figure 1.**
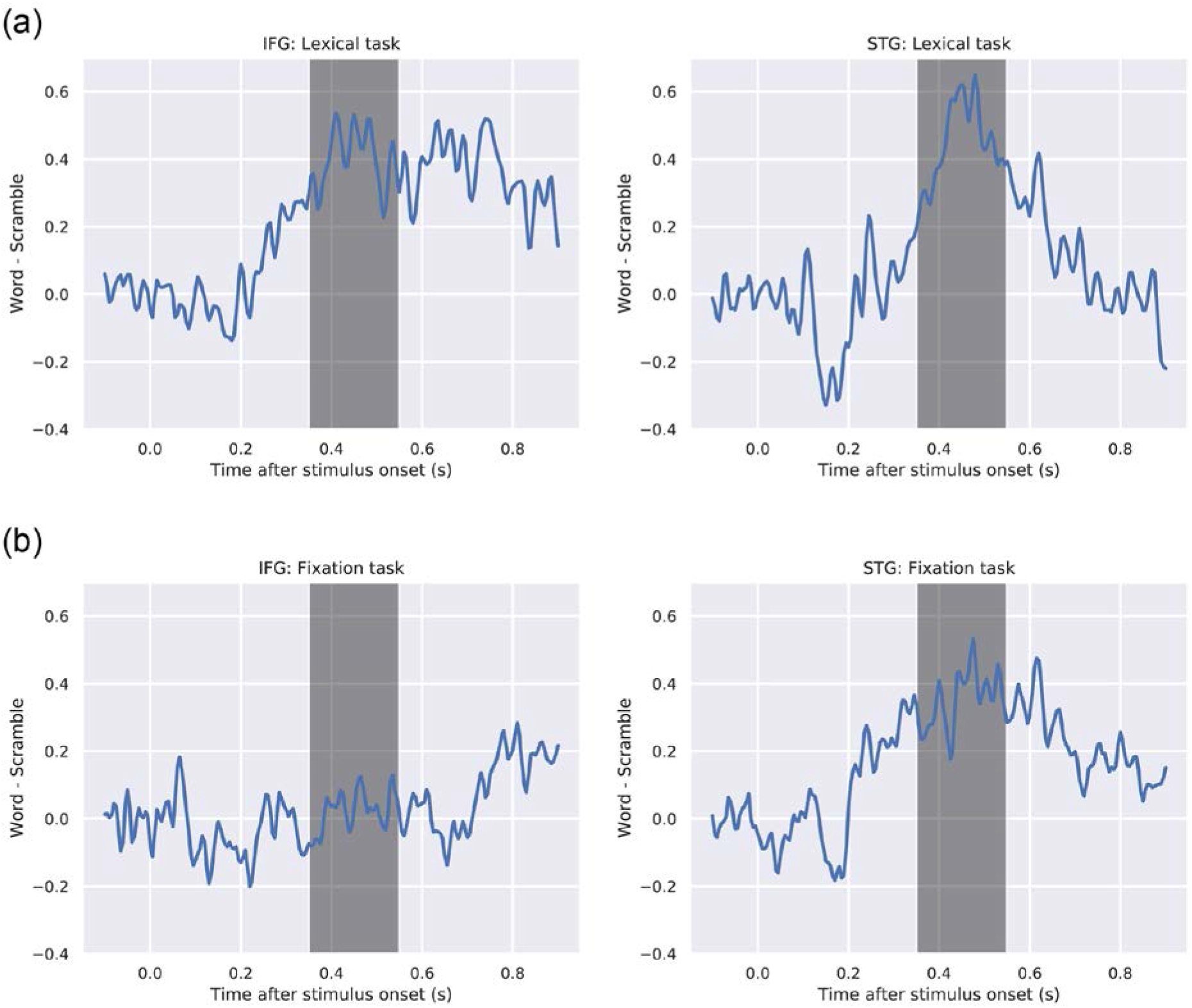
The time course of the word-selective source activity in the lexical (a) and the fixation task (b). The left and the right columns show the word-selective response in the IFG and in the STG, respectively. The shaded region represents the time window of [350, 550] ms and we used this time window to calculated averaged responses at the individual level for the correlation analysis.

**Supplementary Figure 2.**
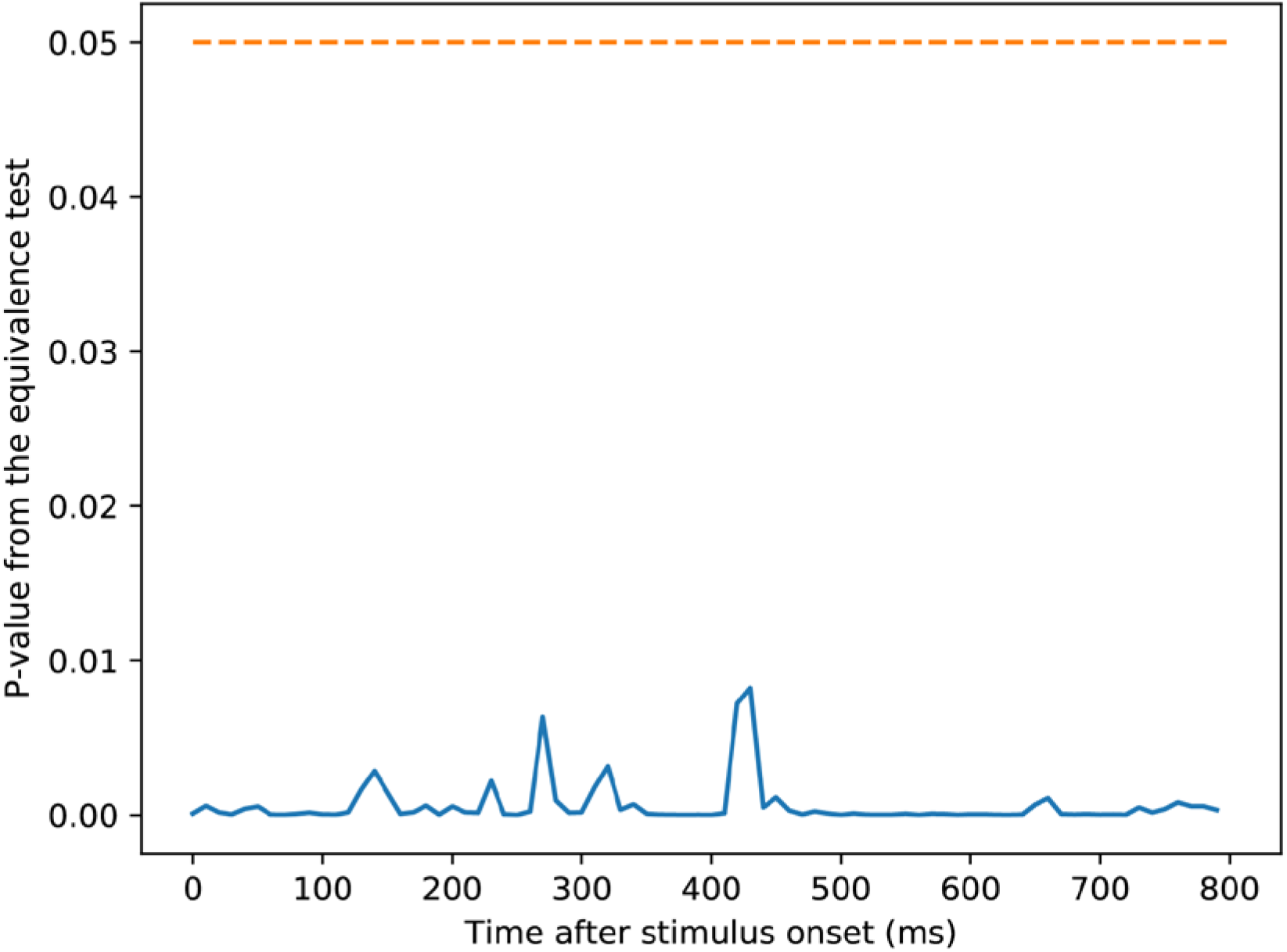
We tested statistical equivalence of word selective responses (words – scramble) in the STG between the lexical and the fixation task. The results suggest that word selective responses in the two tasks were equivalent throughout the epoch. Together with the strong correlation in word selective responses between the two tasks bolsters our claim that the automatic word selective response during the fixation task is similar to the word selective response during active reading.

## References

Bedo, N., Ribary, U., & Ward, L. M. (2014). Fast Dynamics of Cortical Functional and Effective Connectivity during Word Reading. 9(2). https://doi.org/10.1371/journal.pone.0088940

Ben-Shachar, M., Dougherty, R. F., Deutsch, G. K., & Wandell, B. A. (2011). The development of cortical sensitivity to visual word forms. Journal of Cognitive Neuroscience, 23(9), 2387–2399. https://doi.org/10.1162/jocn.2011.21615

Binder, J. R., Frost, J. A., Hammeke, T. A., Cox, R. W., Rao, S. M., & Prieto, T. (1997). Human brain language areas identified by functional magnetic resonance imaging. The Journal of Neuroscience, 17(1), 353–362. https://www.ncbi.nlm.nih.gov/pubmed/8987760

Binder, J. R., Rao, S. M., Hammeke, T. A., Yetkin, F. Z., Jesmanowicz, A., Bandettini, P. A., Wong, E. C., Estkowski, L. D., Goldstein, M. D., & Haughton, V. M. (1994). Functional magnetic resonance imaging of human auditory cortex. Annals of Neurology, 35(6), 662–672. https://doi.org/10.1002/ana.410350606

Blau, V., Reithler, J., Van Atteveldt, N., Seitz, J., Gerretsen, P., Goebel, R., & Blomert, L. (2010). Deviant processing of letters and speech sounds as proximate cause of reading failure: A functional magnetic resonance imaging study of dyslexic children. Brain, 133(3), 868–879. https://doi.org/10.1093/brain/awp308

Blau, V., van Atteveldt, N., Ekkebus, M., Goebel, R., & Blomert, L. (2009). Reduced neural integration of letters and speech sounds links phonological and reading deficits in adult dyslexia. Current Biology, 19(6), 503–508. https://doi.org/10.1016/j.cub.2009.01.065

Blomert, L. (2011). The neural signature of orthographic-phonological binding in successful and failing reading development. Neuroimage, 57(3), 695–703. https://doi.org/10.1016/j.neuroimage.2010.11.003

Brem, S., Bach, S., Kucian, K., Kujala, J. V., Guttorm, T. K., Martin, E., Lyytinen, H., Brandeis, D., & Richardson, U. (2010). Brain sensitivity to print emerges when children learn letter–speech sound correspondences. Proceedings of the National Academy of Sciences, 107(17), 7939–7944. https://doi.org/10.1073/pnas.0904402107

Brunswick, N., McCrory, E., Price, C. J., Frith, C. D., & Frith, U. (1999). Explicit and implicit processing of words and pseudowords by adult developmental dyslexics: A search for Wernicke’s Wortschatz? Brain: A Journal of Neurology, 122 (Pt 1, 1901–1917. https://doi.org/10.1093/brain/122.10.1901

Buracas, G. T., & Boynton, G. M. (2007). The effect of spatial attention on contrast response functions in human visual cortex. The Journal of Neuroscience, 27(1), 93–97. https://doi.org/10.1523/{JNEUROSCI}.3162-06.2007

Caffarra, S., Martin, C. D., Lizarazu, M., Lallier, M., Zarraga, A., Molinaro, N., & Carreiras, M. (2017). Word and object recognition during reading acquisition : MEG evidence. Developmental Cognitive Neuroscience, 24, 21–32. https://doi.org/10.1016/j.dcn.2017.01.002

Chen, Y., Davis, M. H., Pulvermüller, F., & Hauk, O. (2013). Task modulation of brain responses in visual word recognition as studied using EEG/MEG and fMRI. Frontiers in Human Neuroscience, 7(JUN), 376. https://doi.org/10.3389/fnhum.2013.00376

Chen, Yuanyuan, Davis, M. H., Pulvermüller, F., & Hauk, O. (2015). Early Visual Word Processing Is Flexible: Evidence from Spatiotemporal Brain Dynamics. https://doi.org/10.1162/jocn_a_00815

Cornelissen, P. L., Kringelbach, M. L., Ellis, A. W., Whitney, C., Holliday, I. E., & Hansen, P. C. (2009). Activation of the left inferior frontal gyrus in the first 200 ms of reading: evidence from magnetoencephalography ({MEG}). Plos One, 4(4), e5359. https://doi.org/10.1371/journal.pone.0005359

Dale, A. M., Liu, A. K., Fischl, B. R., Buckner, R. L., Belliveau, J. W., Lewine, J. D., & Halgren, E. (2000). Dynamic Statistical Parametric Mapping: Combining fMRI and MEG for High-Resolution Imaging of Cortical Activity. Neuron, 26(1), 55–67. https://doi.org/10.1016/S0896-6273(00)81138-1

Dehaene-lambertz, G., Dehaene, S., & Hertz-pannier, L. (2002). Functional neuroimaging of speech perception. In Infants. Science, 298(December 2002), 2013–2015.

Dehaene, S., Naccache, L., Cohen, L., Bihan, D. L., Mangin, J. F., Poline, J. B., & Rivière, D. (2001). Cerebral mechanisms of word masking and unconscious repetition priming. Nature Neuroscience, 4(7), 752–758. https://doi.org/10.1038/89551

DeWitt, I., & Rauschecker, J. P. (2012). Phoneme and word recognition in the auditory ventral stream. Proceedings of the National Academy of Sciences of the United States of America, 109(8), 2709–2710. https://doi.org/10.1073/pnas.1113427109

Engemann, D. A., & Gramfort, A. (2015). Automated model selection in covariance estimation and spatial whitening of MEG and EEG signals. NeuroImage, 108, 328–342. https://doi.org/10.1016/j.neuroimage.2014.12.040

Fang, F., Boyaci, H., Kersten, D., & Murray, S. O. (2008). Attention-Dependent Representation of a Size Illusion in Human V1. Current Biology, 18(21), 1707–1712. https://doi.org/10.1016/j.cub.2008.09.025

Fischl, B., Sereno, M. I., Tootell, R. B. H., & Dale, A. M. (1999). High-resolution intersubject averaging and a coordinate system for the cortical surface. Human Brain Mapping, 8(4), 272–284. https://doi.org/10.1002/(SICI)1097-0193(1999)8:4<272::AID-HBM10>3.0.CO;2-4

Gramfort, A., Luessi, M., Larson, E., Engemann, D. A., Strohmeier, D., Brodbeck, C., Goj, R., Jas, M., Brooks, T., Parkkonen, L., & Hämäläinen, M. (2013). MEG and EEG data analysis with MNE-Python. Frontiers in Neuroscience, 7 DEC. https://doi.org/10.3389/fnins.2013.00267

Gross, J., Baillet, S., Barnes, G. R., Henson, R. N., Hillebrand, A., Jensen, O., Jerbi, K., Litvak, V., Maess, B., Oostenveld, R., Parkkonen, L., Taylor, J. R., van Wassenhove, V., Wibral, M., & Schoffelen, J. M. (2013). Good practice for conducting and reporting MEG research. NeuroImage, 65, 349–363. https://doi.org/10.1016/j.neuroimage.2012.10.001

Harm, M W, & Seidenberg, M. S. (1999). Phonology, reading acquisition, and dyslexia: insights from connectionist models. Psychological Review, 106(3), 491–528. https://doi.org/10.1037/0033-{295X}.106.3.491

Harm, Michael W, & Seidenberg, M. S. (2004). Computing the meanings of words in reading: cooperative division of labor between visual and phonological processes. Psychological Review, 111(3), 662–720. https://doi.org/10.1037/0033-{295X}.111.3.662

Hebb, D. O. (1949). The Organization of Behavior. A Neuropsychological Theory. John Wiley and Sons, Inc. https://psycnet.apa.org/record/1950-02200-000

Heilbron, M., Richter, D., Ekman, M., Hagoort, P., & de Lange, F. P. (2020). Word contexts enhance the neural representation of individual letters in early visual cortex. Nature Communications, 11(1), 1–11. https://doi.org/10.1038/s41467-019-13996-4

Helenius, P., Salmelin, R., Service, E., & Connolly, J. F. (1998). Distinct time courses of word and context comprehension in the left temporal cortex. Brain, 121(6), 1133–1142. https://doi.org/10.1093/brain/121.6.1133

Kay, K. N., & Yeatman, J. D. (2017). Bottom-up and top-down computations in word- and face-selective cortex. ELife, 6. https://doi.org/10.7554/{eLife}.22341

Klein, M., Grainger, J., Wheat, K. L., Millman, R. E., Simpson, M. I. G., Hansen, P. C., & Cornelissen, P. L. (2015). Early Activity in Broca’s Area during Reading Reflects Fast Access to Articulatory Codes from Print. Cerebral Cortex, 25(7), 1715–1723. https://doi.org/10.1093/cercor/bht350

Kriegeskorte, N., Simmons, W. K., Bellgowan, P. S., & Baker, C. I. (2009). Circular analysis in systems neuroscience: The dangers of double dipping. Nature Neuroscience, 12(5), 535–540. https://doi.org/10.1038/nn.2303

Lerma-Usabiaga, G., Carreiras, M., & Paz-Alonso, P. M. (2018). Converging evidence for functional and structural segregation within the left ventral occipitotemporal cortex in reading. Proceedings of the National Academy of Sciences. https://doi.org/10.1073/pnas.1803003115

Luck, S. J., Vogel, E. K., & Shapiro, K. L. (1996). Word meanings can be accessed but not reported during the attentional blink. In Nature (Vol. 383, Issue 6601, pp. 616–618). https://doi.org/10.1038/383616a0

Mano, Q. R., Humphries, C., Desai, R. H., Seidenberg, M. S., Osmon, D. C., Stengel, B. C., & Binder, J. R. (2013). The role of left occipitotemporal cortex in reading: Reconciling stimulus, task, and lexicality effects. Cerebral Cortex, 23(4), 988–1001. https://doi.org/10.1093/cercor/bhs093

Maris, E., & Oostenveld, R. (2007). Nonparametric statistical testing of EEG- and MEG-data. Journal of Neuroscience Methods, 164(1), 177–190. https://doi.org/10.1016/j.jneumeth.2007.03.024

McCandliss, B. D., Cohen, L., & Dehaene, S. (2003). The visual word form area: Expertise for reading in the fusiform gyrus. Trends in Cognitive Sciences, 7(7), 293–299. https://doi.org/10.1016/S1364-6613(03)00134-7

Mesgarani, N., Cheung, C., Johnson, K., & Chang, E. F. (2014). Phonetic feature encoding in human superior temporal gyrus. TL - 343. Science (New York, N.Y.), 343 VN-(6174), 1006–1010. https://doi.org/10.1126/science.1245994

Norton, E. S., & Wolf, M. (2012). Rapid Automatized Naming (RAN) and Reading Fluency: Implications for Understanding and Treatment of Reading Disabilities. Annual Review of Psychology, 63(1), 427–452. https://doi.org/10.1146/annurev-psych-120710-100431

Pattamadilok, C., Chanoine, V., Pallier, C., Anton, J.-L., Nazarian, B., Belin, P., & Ziegler, J. C. (2017). Automaticity of phonological and semantic processing during visual word recognition. NeuroImage, 149(February), 244–255. https://doi.org/10.1016/j.neuroimage.2017.02.003

Paulesu, E. (2001). Dyslexia: Cultural Diversity and Biological Unity. Science, 291(5511), 2165–2167. https://doi.org/10.1126/science.1057179

Paulesu, E., Frith, C. D., & Frackowiak, R. S. (1993). The neural correlates of the verbal component of working memory. Nature, 362(6418), 342–345. https://doi.org/10.1038/362342a0

Perfetti, C. A., & Bell, L. (1991). Phonemic activation during the first 40 ms of word identification: Evidence from backward masking and priming. Journal of Memory and Language, 30(4), 473–485. https://doi.org/10.1016/0749-596X(91)90017-E

Perfetti, C. A., Bell, L. C., & Delaney, S. M. (1988). Automatic (prelexical) phonetic activation in silent word reading: Evidence from backward masking. Journal of Memory and Language, 27(1), 59–70. https://doi.org/10.1016/0749-{596X}(88)90048-4

Pernet, C. R., Wilcox, R., & Rousselet, G. A. (2013). Robust correlation analyses: False positive and power validation using a new open source matlab toolbox. Frontiers in Psychology, 3(JAN), 1–18. https://doi.org/10.3389/fpsyg.2012.00606

Preston, J. L., Molfese, P. J., Frost, S. J., Mencl, W. E., Fulbright, R. K., Hoeft, F., Landi, N., Shankweiler, D., & Pugh, K. R. (2016). Print-Speech Convergence Predicts Future Reading Outcomes in Early Readers. Psychological Science, 27(1), 75–84. https://doi.org/10.1177/0956797615611921

Price, C. J. (2012). A review and synthesis of the first 20 years of {PET} and {fMRI} studies of heard speech, spoken language and reading. Neuroimage, 62(2), 816–847. https://doi.org/10.1016/j.neuroimage.2012.04.062

Price, C. J., Wise, R. J. S., & Frackowiak, R. S. J. (1996). Demonstrating the implicit processing of visually presented words and pseudowords. Cerebral Cortex, 6(1), 62–70. https://doi.org/10.1093/cercor/6.1.62

Pugh, K R, Frost, S. J., Sandak, R., Landi, N., Moore, D., Porta, G. D., Rueckl, J. G., & Mencl, W. E. (2010). Mapping the word reading circuitry in skilled and disabled readers. In P. Cornelissen, P. Hansen, M. Kringelbach, & K. Pugh (Eds.), The Neural Basis of Reading. Oxford University Press.

Pugh, K R, Mencl, W. E., Jenner, A. R., Katz, L., Frost, S. J., Lee, J. R., Shaywitz, S. E., & Shaywitz, B. A. (2001). Neurobiological studies of reading and reading disability. Journal of Communication Disorders, 34(6), 479–492. https://doi.org/10.1016/s0021-9924(01)00060-0

Pugh, Kenneth R., Frost, S. J., Sandak, R., Landi, N., Moore, D., Porta, G. D., Rueckl, J. G., & Mencl, W. E. (2010). Mapping the Word Reading Circuitry in Skilled and Disabled Readers. In P. Cornelissen, P. Hansen, M. Kringelbach, & K. Pugh (Eds.), The Neural Basis of Reading (pp. 281–305). Oxford University Press.

Raij, T., Uutela, K., & Hari, R. (2000). Audiovisual integration of letters in the human brain. Neuron, 28(2), 617–625. https://doi.org/10.1016/S0896-6273(00)00138-0

Rauschecker, A. M., Bowen, R. F., Parvizi, J., & Wandell, B. A. (2012). Position sensitivity in the visual word form area. Proceedings of the National Academy of Sciences, 109(24), E1568–E1577. https://doi.org/10.1073/pnas.1121304109

Rauschecker, Andreas M., Bowen, R. F., Perry, L. M., Kevan, A. M., Dougherty, R. F., & Wandell, B. A. (2011). Visual feature-tolerance in the reading network. Neuron, 71(5), 941–953. https://doi.org/10.1016/j.neuron.2011.06.036

Reicher, G. M. (1969). Perceptual recognition as a function of meaningfulness of stimulus material. Journal of Experimental Psychology, 81(2), 275–280. https://doi.org/10.1037/h0027768

Saffran, J R, Senghas, A., & Trueswell, J. C. (2001). The acquisition of language by children. Proceedings of the National Academy of Sciences of the United States of America, 98(23), 12874–12875. https://doi.org/10.1073/pnas.231498898

Saffran, Jenny R., Aslin, R. N., & Newport, E. L. (1996). Statistical learning by 8-month-old infants. Science, 274(5294), 1926–1928. https://doi.org/10.1126/science.274.5294.1926

Sahin, N. T., Pinker, S., Cash, S. S., Schomer, D., & Halgren, E. (2009). Sequential processing of lexical, grammatical, and phonological information within Broca’s area. Science, 326(5951), 445–449. https://doi.org/10.1126/science.1174481

Seidenberg, M. S., & McClelland, J. L. (1989). A distributed, developmental model of word recognition and naming. Psychological Review, 96(4), 523–568. https://doi.org/10.1037/0033-295X.96.4.523

Strijkers, K., Bertrand, D., & Grainger, J. (2015). Seeing the same words differently: The time course of automaticity and top-down intention in reading. Journal of Cognitive Neuroscience, 27(8), 1542–1551. https://doi.org/10.1162/jocn_a_00797

Stroop, J. R. (1935). Studies of interference in serial verbal reactions. Journal of Experimental Psychology, 18(6), 643–662. https://doi.org/10.1037/h0054651

Tarkiainen, A., Helenius, P., Hansen, P. C., Cornelissen, P. L., & Salmelin, R. (1999). Dynamics of letter string perception in the human occipitotemporal cortex. Brain, 122(11), 2119–2131. https://doi.org/10.1093/brain/122.11.2119

Taulu, S., Simola, J., & Kajola, M. (2005). Applications of the signal space separation method. IEEE Transactions on Signal Processing, 53(9), 3359–3372. https://doi.org/10.1109/TSP.2005.853302

Taylor, J. S. H., Davis, M. H., & Rastle, K. (2019). Mapping visual symbols onto spoken language along the ventral visual stream. Proceedings of the National Academy of Sciences, 201818575. https://doi.org/10.1073/pnas.1818575116

Turkeltaub, P. E., Gareau, L., Flowers, D. L., Zeffiro, T. A., & Eden, G. F. (2003). Development of neural mechanisms for reading. Nature Neuroscience, 6(7), 767–773. https://doi.org/10.1038/nn1065

van Atteveldt, N., Formisano, E., Goebel, R., & Blomert, L. (2004). Integration of letters and speech sounds in the human brain. Neuron, 43(2), 271–282. https://doi.org/10.1016/j.neuron.2004.06.025

Van Orden, G. C., & Goldinger, S. D. (1994). Interdependence of Form and Function in Cognitive Systems Explains Perception of Printed Words. Journal of Experimental Psychology: Human Perception and Performance, 20(6), 1269–1291. https://doi.org/10.1037/0096-1523.20.6.1269

Wandell, B. A., Rauschecker, A. M., & Yeatman, J. D. (2012). Learning to See Words. Annual Review of Psychology, 63(1), 31–53. https://doi.org/10.1146/annurev-psych-120710-100434

Warrington, E. K., Logue, V., & Pratt, R. T. (1971). The anatomical localisation of selective impairment of auditory verbal short-term memory. Neuropsychologia, 9(4), 377–387. https://doi.org/10.1016/0028-3932(71)90002-9

Wheat, K. L., Cornelissen, P. L., Frost, S. J., & Hansen, P. C. (2010). During visual word recognition, phonology is accessed within 100 ms and may be mediated by a speech production code: evidence from magnetoencephalography. The Journal of Neuroscience, 30(15), 5229–5233. https://doi.org/10.1523/{JNEUROSCI}.4448-09.2010

Wheeler, D. D. (1970). Processes in word recognition. Cognitive Psychology, 1(1), 59–85. https://doi.org/10.1016/0010-0285(70)90005-8

Wilson, S. M., Saygin, A. P., Sereno, M. I., & Iacoboni, M. (2004). Listening to speech activates motor areas involved in speech production. Nature Neuroscience, 7(7), 701–702. https://doi.org/10.1038/nn1263

Wise, R., Chollet, F., Hadar, U., Friston, K., Hoffner, E., & Frackowiak, R. (1991). Distribution of cortical neural networks involved in word comprehension and word retrieval. Brain: A Journal of Neurology, 114 (Pt 4, 1803–1817. https://doi.org/10.1093/brain/114.4.1803

Wolf, M. (2018). Reader, come home : the reading brain in a digital world. Harper.

Yi, H. G., Leonard, M. K., & Chang, E. F. (2019). The Encoding of Speech Sounds in the Superior Temporal Gyrus. In Neuron (Vol. 102, Issue 6, pp. 1096–1110). Cell Press. https://doi.org/10.1016/j.neuron.2019.04.023

